# Spatial Venomics - Cobra Venom System Reveals Spatial Differentiation of Snake Toxins by Mass Spectrometry Imaging

**DOI:** 10.1101/2022.01.31.478453

**Authors:** Benjamin-Florian Hempel, Maik Damm, Daniel Petras, Taline D. Kazandjian, Claudia A. Szentiks, Guido Fritsch, Grit Nebrich, Nicholas R. Casewell, Oliver Klein, Roderich D. Süssmuth

**Affiliations:** BIH Center for Regenerative Therapies BCRT, Charité - Universitätsmedizin Berlin, 13353 Berlin, Germany; Institut für Chemie, Technische Universität Berlin, 10623 Berlin, Germany; CMFI Cluster of Excellence, Interfakultäres Institut für Mikrobiologie und Infektionsmedizin Tübingen, Universität Tübingen, 72076 Tübingen, Germany; Centre for Snakebite Research & Interventions, Liverpool School of Tropical Medicine, Liverpool, L3 5QA, United Kingdom; Department of Wildlife Diseases and Reproduction Managment, Leibniz-Institut für Zoo- und Wildtierforschung (IZW) im Forschungsverbund Berlin e.V., 10315 Berlin, Germany

**Keywords:** venom gland morphology, mass spectrometry imaging, spatial venomics, proteomics, venom heterogeneity

## Abstract

Among venomous animals, toxic secretions have evolved as biochemical weapons associated with various highly specialized delivery systems on many occasions. Despite extensive research, there is still limited knowledge of the functional biology of most animal toxins, including their venom production and storage, as well as the morphological structures within sophisticated venom producing tissues that might underpin venom modulation. Here we report on the spatial exploration of a snake venom gland system by matrix-assisted laser desorption/ionization mass spectrometry imaging (MALDI-MSI), in combination with standard proteotranscriptomic approaches, to enable in situ toxin mapping in spatial intensity maps across a venom gland sourced from the Egyptian cobra (*Naja haje*). MALDI-MSI toxin visualization on the elapid venom gland reveals high spatial heterogeneity of different toxin classes at the proteoform level, which may be the result of physiological constraints on venom production and/or storage that reflects the potential for venom modulation under diverse stimuli.

## Introduction

Venoms are sophisticated and complex mixtures consisting of low molecular weight compounds, peptides and proteins, which have evolved for use as defensive and/or foraging adaptations in various animal lineages.^[1,2]^ Venom research is the focus of various scientific disciplines due to their great medical importance, i.e. the generation of effective immunological therapies against envenoming and the discovery of novel drugs from venoms.^[3,4]^ The structural and functional diversity of venom systems has been triggered by a variety of adaptive evolutionary processes, such as convergence^[1]^, co-option of single copy genes^[5]^, gene duplication^[6]^, accelerated rates of molecular evolution^[7]^, and protein neofunctionalization^[8]^. Thus, collectively venoms have proven to be valuable models for examining the origins of adaptations and the link between genotype and functional phenotypes.

Among venomous animals, serpents have received the most scientific attention due to their frequently lethal encounters with human.^[4]^ Snake venoms are among the most lethal bioweapons in nature that are produced, stored and released by a sophisticated venom delivery system in the upper jaw.^[9]^ This secretion, produced in specialized tissues and embedded in the venom gland apparatus, causes a cascade of physiological and biochemical perturbations once delivered into the target (e.g. prey, predator or aggressor).^[1]^ The complex arsenal found in venomous snakes is an unique characteristic and knowledge of its molecular composition has aid to categorize major serpent families, as venom toxin variation is ubiquitous across multiple taxonomic levels.^[10,11]^ A typical feature of elapid venoms is the abundant presence of the non-enzymatic three-finger toxins (3FTx’s) beside a less diverse number of other toxin families.^[12,13]^ The 3FTx’s are a broad toxin superfamily that interferes with a variety of biological functions of the prey/victim by interacting with various molecular targets, including receptors, ion channel proteins and enzymes.^[14,15]^ Venoms are energetically expensive commodities and the degree of toxin diversity in relation to the morphological attributes of the venom system substantially dependent on the degree to which an animal relies on biochemical versus physical means for overpowering prey.^[16]^ Although mass spectrometry-based (MS) venom proteomics and sequencing-based methods facilitate detailed insight into venom diversity, classical proteomics and transcriptomics analyses using venom secretions or even tissue homogenates are not able to correlate and spatially resolve single toxins within cellular environments.^[17]^

Matrix-assisted laser desorption/ionization mass spectrometry imaging (MALDI-MSI) can act at the interface of venom diversity and spatial localization to bypass existing limitations of classical venomics approaches. It allows unsupervised and simultaneous analyses of molecules (e.g., metabolites, proteins, peptides, lipids and glycans) in situ on a single tissue section, preserving their spatial coordinates and generating a molecular intensity map reflecting the relative molecule abundance.^[17,18]^ The continual technical improvements of MALDI-MSI has raised its suitability and applicability to become a valuable tool for use in the field of venomics. Hence, it facilitates mapping of individual toxins on-tissue and provides deeper insights into the spatial venom distribution as well as heterogeneity and its effects on venom regulation or modulation.^[19]^ The localization of different venom toxins within the context of morphological structures has previously been described by MALDI-MSI in a multidimensional manner.^[20–22]^ Ghezellou *et al*.^[23]^ used atmospheric-pressure MALDI (AP-MALDI) to confirm the presence of various lipids within the saw-scaled viper (*Echis carinatus sochureki*) venom system. However, previous MSI experiments are restricted in their venom identification, which allows intact profiling of selected toxin classes and excludes the identification of higher molecular weight toxins, like snake venom serine proteases (svSP), snake venom metalloproteases (svMP), or L-amino acid oxidases (LAAO).

Here we demonstrate the feasibility and potential of MALDI-MSI, in combination with a global venom analysis via a proteotranscriptomics approach, as the basis for the spatial identification of different toxins associated with the venom system of the Egyptian cobra (*Naja haje*, formerly *N. h. legionis*, locale: Morocco).^[19]^ Therefore, tissue sections of the elapid venom gland were enzymatically digested on-tissue to allow the identification of small and high molecular weight toxin classes in high spatial resolution. In combination with an analysis of the morphological structures and histological features of the elapid venom gland, we depict a holistic venom overview indicating that venom production and storage is a highly complex process. Our spatial venom analysis helps to place the simple and centralized elapid venom system in an evolutionary context, gaining insight into the complexity of toxin compartmentalization, which may be the result of physiological constraints on venom production and/or storage, or reflect the potential for venom modulation under different stimuli.

## Results and Discussion

### Morphology of the Egyptian cobra (*N. haje*) venom gland

To visualize the cobra venom production and storage system and to obtain a better understanding of the physiological structures and relative position of an elapid venom gland, we performed three-dimensional computed tomography (3D-CT) and histochemical staining (**Figure 1**).

**Figure 1.**
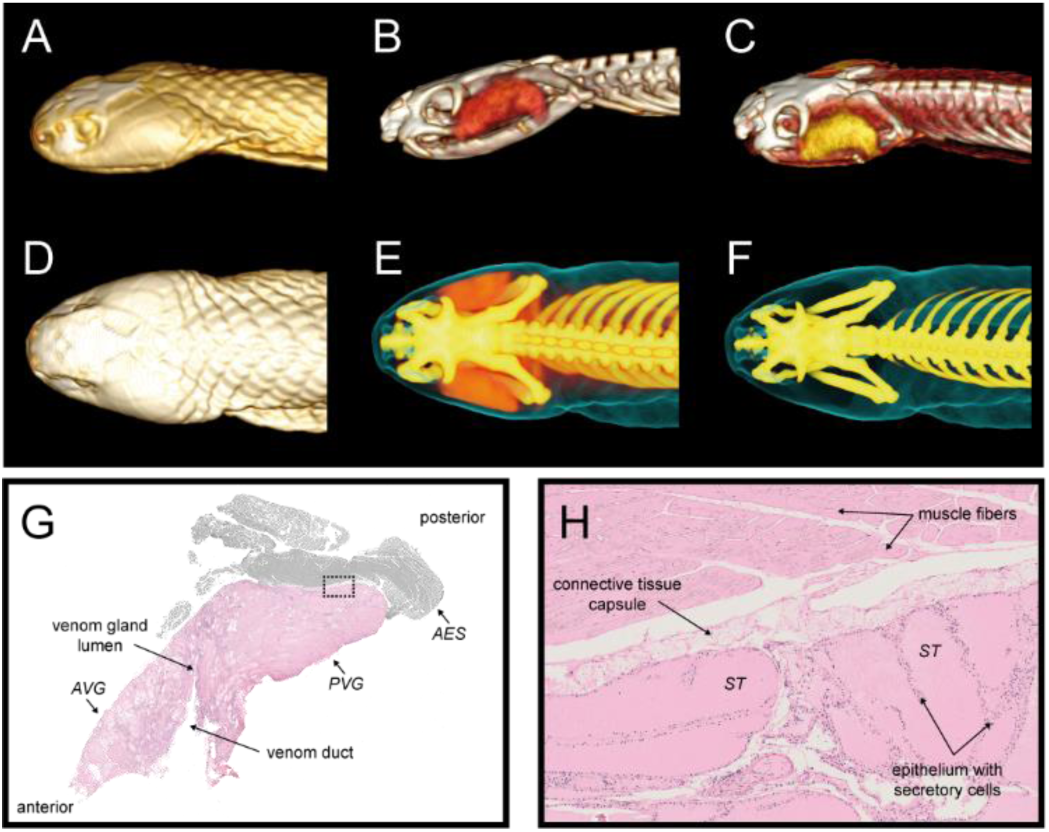
Morphology, topography and histochemistry of the venom delivery system of *Naja haje*. (**A-F**) Three-dimensional computer tomography (3D-CT) reconstruction showing the relative position (lateral and top view) of the cobra head (**A, D**), morphological location of the venom production system (**B, E**), and skeleton with and without venom gland system (**C, F**). (**G, H**) Longitudinal section and histochemical staining by eosin and hematoxylin (H/E) of the *N. haje* venom gland. (**G**) The histochemical staining of a sagittal section in posteroanterior direction within the venom gland system. It reveals the *adductor externus super?cialis* (AES), dorsal and caudal attached to the venom system. Both delimited by a thick connective tissue capsule (*capsula fibrosa*). The venom duct, associated to a narrow lumen, divides the inner gland structure in an anterior (AVG) and a posterior venom gland (PVG) regime. (**H**) The venom duct region is formed by a high proportion of connective tissue and some strong elongated follicle, so-called secretory tubules (ST), RVG and CVG are composed of large and oval STs and thin connective tissue (zoom-in for rectangle in G).

Morphological and phylogenetic biology studies on the serous-secreting venom glands in different caenophidian snake families suggest a single origin at the base of the colubroid radiation between 60-80 Mya.^[10]^ Since this early origin, associated morphological characteristics of the venom system have evolved independently on multiple occasions in different caenophidian taxons, resulting in particular structural and topological characteristics, including fangs and glandular system.^[24–26]^ Advanced snakes have centralized venom systems to store their ready-to-use venom cocktail produced within. The simple glandular system might permit quantitative regulation of secreted venom depending on the ecological stimuli a snake is presented with. The 3D-CT reconstruction gave insights into the anatomy reflected by different layers of depth and revealed extensions of the venom glands and their general architecture (**Figure 1A-F**).

In the first level, a typical cobra head is shown covered with lateral hoods and small, round pupils in lateral or top view, respectively (**Figure 1A** and **D**). Furthermore, the detailed topographical location of the venom gland system and skeleton with and without surrounded muscular tissue in lateral (**Figure 1B** and **C**) or top view (**Figure 1E** and **F**) is shown, respectively. The slender anterior ends of the elapid venom glands are positioned postorbital associated on each side of the head with specialized venom-conducting fangs located on the maxilla, as in all other advanced snakes (**Figure 1B and E**).^[9]^ The basic structure shows an oval shape and is more compact in contrast to large and triangular-shaped venom glands of viperids (**Figure 1C and E**).^[27]^

The main function of the venom glands is the production and storage of a specialized toxic secretion prior to its delivery via the fangs into prey or aggressors.^[27]^ Its postorbital position and the roughly oval shape is consistent with other previously described studies for closely related species of *Elapidae* and is a typical representative of a centralized and simple venom system.^[24]^ However, it is worth mentioning that venom glands differ considerably in form and internal fine structure amongst taxa, sometimes even between species within the same genus.^[26]^

The histochemical section in sagittal orientation reveals the closely associated *adductor externus super?cialis* (AES), dorsally and caudally attached to elapid venom glands, and responsible for venom release after compression (**Figure 1G**).^[24,26,28]^ Furthermore, this compressor muscle is attached to a thick connective tissue capsule (*capsula fibrosa*) that encloses the venom gland and clearly delimits it from the AES (**Figure 1H**).^[24,29]^ The roughly centered venom duct, which is directly associated to a narrow lumen above, runs in a posteroanterior direction and divides the inner gland structure in an anterior (AVG) and a posterior venom gland (PVG) regime (**Fig. 1G**). While the venom duct region is formed by a high proportion of connective tissue and some strong elongated follicle, so called secretory tubules (ST), AVG and PVG are mainly composed of large and oval STs (∼260 µm ± 70 µm in diameter) surrounded by thin connective tissue (**Figure 1H**).^[24]^ Due to the relatively small size, the venom gland lumen is unsuitable for long-term storage of venom secretion. In consequence, the venom is stored within the STs rather than the lumen. The outer surface of the STs are formed by a single layer of epithelium with secretory cells, which enclose the colloid-like lumen and shapes the venom gland fine structure analog to other oral secretory systems, also known for secretion and long term storage (**Figure 1H**).^[28,30]^ This underlying morphological data of longitudinal venom gland sections can help to put the localization and visualization of numerous venom peptides into a morphological context.

### Proteotranscriptomics of Egyptian cobra venom

To examine the secretory output of the elapid venom gland, we utilized a set of transcriptomics and complementary bottom-up and top-down proteomics data, thereby providing a holistic overview of venom composition (**Table S1-5**).

The *N. haje* venom gland transcriptome resulted in 4,418 assembled contigs, of which 58 exhibited gene annotations relating to 18 venom toxin families (**Figure 2A** and **Table S1**).^[31]^ Of these contigs, six were full-length toxin sequences that encoded toxin isoforms relating to 3FTx family. This toxin family exhibited the highest expression levels by far of all toxin families identified, in combination accounting for >86% of the total toxin expression. In particular, long neurotoxins (lNTx) exhibited the highest expression level (41%) within the 3FTx family, followed by short neurotoxins (sNTx, 39%), weak neurotoxins (wNTx, 5%) and cytotoxins (CTx, 2%). The Kunitz-type inhibitor (KUN), snake venom metalloproteinases (svMP) and serine proteases (svSP) were the second most abundant toxin families representing each ∼3% of the toxin gene expression. Lower expression levels were identified for cysteine-rich secretory protein (CRISP), phospholipase A_2_ inhibitor (PLA_2_-i), cobra venom factor (CVF), venom nerve growth factor (NGF), L-amino acid oxidase (LAAO), 5’-nucleotidase (5NUC), and phosphodieseterase (PDE), which account each ≥1% of total toxin expression (**Figure 2A**; secondary). A number of other potential venom toxin families (**Figure 2A;** minor) were identified with very low expression levels (in total <1%) (**Table S1**).

**Figure 2.**
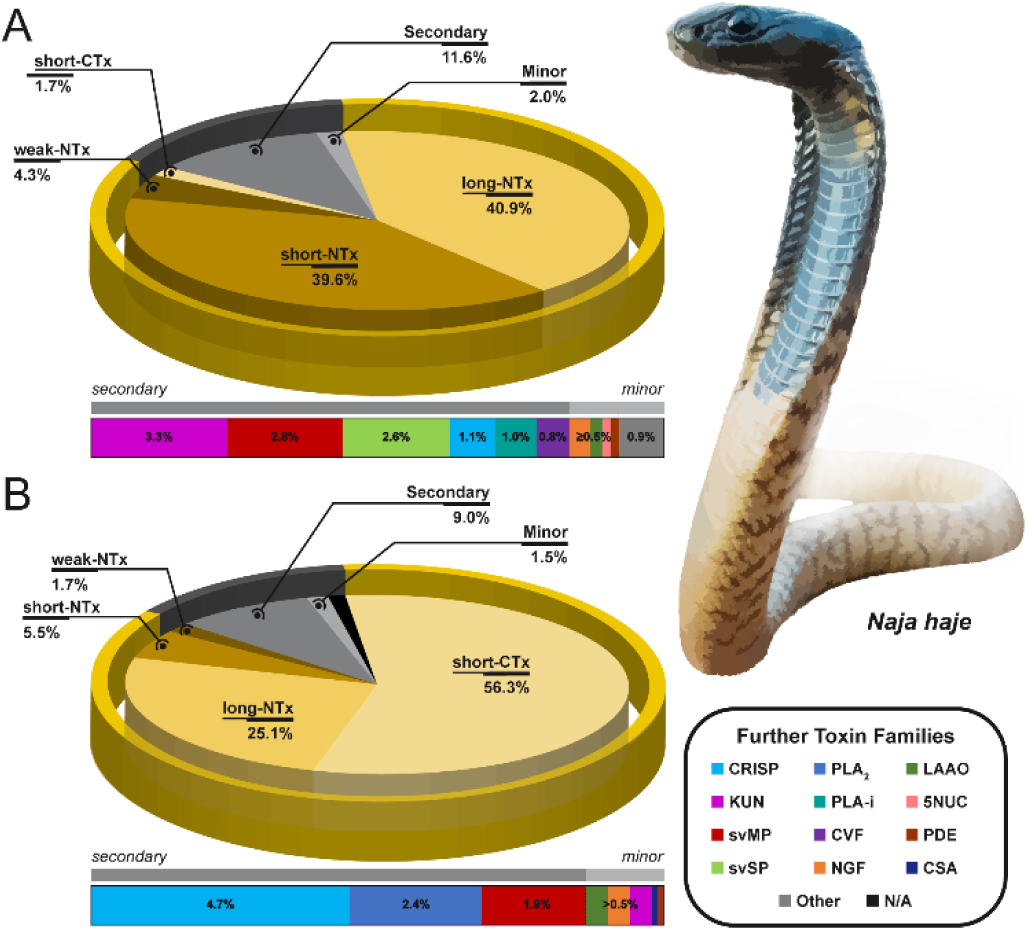
Proteotranscriptomics analysis of the Egyptian cobra (*Naja haje*). The relative gene expression levels (**A**) and protein abundance (**B**) of toxin families in percentage values identified in the *N. haje* venom gland transcriptome and venom proteome, respectively (**Table S1** and **S2**). (**A**) The relative gene expression levels (transcriptome) of the major 3FTx toxin family and subgroups, secondary (>5%) as well as minor (>0.5%) toxin families represented by a pie chart. The corresponding bar chart reflects the relative expression levels of the secondary and minor categorized toxin families, which in combination account for 13.6% of residual toxins encoded in the venom gland. (**B**) The relative venom protein abundance (proteome) of the major 3FTx toxin family and subgroups, secondary (>5%) as well as minor (>0.5%) toxin families represented by a pie chart. The corresponding bar chart reflects the relative protein abundance of the secondary and minor categorized toxin families, which in combination account for 10.5% of residual toxins present in the venom proteome. The color code (bottom right) assigns to the various toxin superfamilies.

Next, we applied shotgun proteomics to broadly characterize the composition of secreted venom, including the qualitative assessments of main toxin families and the generation of a peptide reference database for later annotation of MALDI-MSI data (**Table S2**). Therefore, we pursued a dual strategy by applying in-solution digestion of crude venom combined with an on-top tissue digestion of an adjacent section. Overall, the combined shotgun approach resulted in 648 peptide spectrum matches (PSM) at a false discovery rate (FDR) of 0.1% to our species-specific assembled transcriptome database, extended by various *Naja* species^[31]^ and the NCBI protein database (taxid: 8602). The toxin-encoding contigs covered 16 different toxin classes. We identified a wide range of typical elapid toxin families, such as 3FTx, CRISP, PLA_2_, svSP, svMP, LAAO, NGF, KUN, PDE, 5NUC, CVF, and other lowly abundant venom toxins (**Table S2**). Further, all sub-groups of the 3FTx superfamily could be assigned.^[14]^ Interestingly, related elapid species showed the presence of other less abundant toxin classes such as cystatin (CYS), acetylcholinesterase (AChE), and natriuretic peptides (NP) that are fully absent in the venom proteome described here.^[32]^ On the other hand, we identified several peptide masses belonging to PLA_2_-i and a muscarinic toxin-like protein (MTLP).^[12]^

In addition, we performed a semi-quantitative bottom-up analysis by reversed phase-HPLC (**Figure S2A**) and subsequent SDS-PAGE (**Figure S2B**) separation, as already described in detail elsewhere.^[33]^ The decomplexation of the *N. haje* venom resulted in 44 characteristic fractions, which separated into 151 protein bands covering a mass range of 5-100 kDa. Based on the comprehensive transcriptome database, we annotated respective MS/MS data to assign venom toxin families (**Figure 2B**). The 595 annotated PSMs resulted in the identification of various proteins from 12 different toxin families. The most abundant toxin family is represented by 3FTx (88%), with its subgroups: sNTx (6%), lNTx (25%), wNTx (2%) and CTx (56%). Secondary toxin families including CRISP (5%), PLA_2_ (3%), and svMP (2%) are followed by low abundant toxin families, such LAAO (0.5%), KUN (0.4%), NGF (0.4%), cobra serum albumin (CSA; 0.1%), and PDE (0.1%). Further, PLA_2_-i, 5NUC and CVF were only present in trace amounts, alongside a minor amount of unannotated proteins (N/A; 1.0%) (**Table S3** and **S4**). Employing both proteomics and transcriptomics, rather than only one of these techniques, improved the generation of a comprehensive and accurate venom database. When comparing the abundance of venom toxins (**Figure 2B**) with transcriptomic predictions of expression levels (**Figure 2A**), we observed an overall positive correlation, but noted some considerable differences, particular for the subgroups of the 3FTx family. The observed discrepancies in proteomic abundance and transcriptomic expression levels (e.g. sNTx and CTx; **Figure 2**) might be influenced by a number of biological and experimental factors, as previously outlined.^[34,35]^ Perhaps the most pressing reason is that we compared toxin transcription levels from a Ugandan individual with toxin abundance calculations from a Moroccan specimen.^[35,36]^ Thus, while it is possible that these differences are predominantly due to the abovementioned regulatory processes, it seems likely that intra-species variations influence our proteotranscriptomics results (**Figure 2**).^[13]^

Top-down mass profiling by high-resolution electrospray ionization (HR-ESI) MS of the native Moroccan cobra venom generated an overview of 64 unique intact toxin isoforms, including low abundant compounds and small peptides (**Figure S3** and **Table S3**). The intact mass profile (IMP) is dominated by 43 molecular masses in the range of 6,377-7,897 Da, which reflects a considerable variety of 3FTx, as shown in detail by the bottom-up approach. Within the 3FTx family, we identified the major subclasses, such as CTx with ‘Cytotoxin 2’ (6,845.25 Da; CTx-2), followed by ‘Cytotoxin 5’ (6,761.21 Da; CTx-5), ‘Cytotoxin 6’ (6,813.30 Da; CTx-6) and ‘Cytotoxin 11’ (6,829.55 Da; CTx-11) and the neurotoxins (NTx) that are dominated by ‘long Neurotoxin 1’ (7,896.50 Da; lNTx-1). Furthermore, KUN were detected in three fractions, showing only a marginal part with two different proteoforms (6,377 Da and 6,392 Da). CRISP could be associated with nine different proteoform masses between 23-25 kDa. The largest observed intact mass of 49 kDa was identified as a single svMP proteoform, the main representative of proteases in this venom.

### Colocalization of molecular features to morphological structures

In order to prove complex morphological structures by discriminative molecular pattern and identify subregions across a venom system section of the Egyptian cobra (*N. haje*), we performed unsupervised multivariate analyses of acquired *m/z* values (peak features) without any need to extract or label samples. (**Figure 3**).

**Figure 3.**
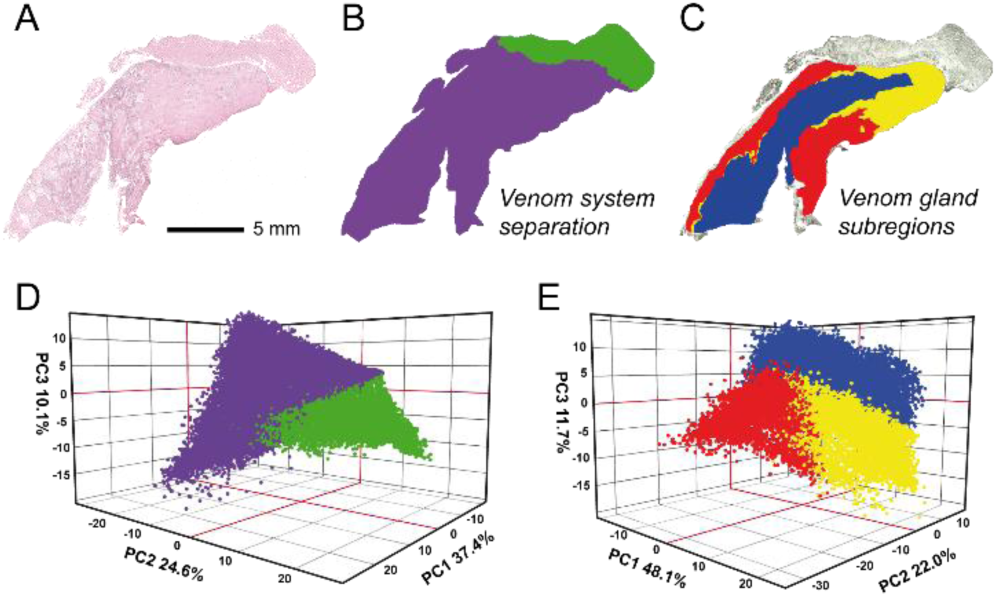
Multivariate statistical analyses of the venom gland system. Spatial segmentation generated by bisecting k-means clustering represents different hierarchical clusters (indicated by different colors) based on discriminative *m/z* values (peak features). (**A**) The histochemical staining with hematoxylin and eosin (H&E) for tissue section orientation in segmentation maps of MALDI-MSI analysis. (**B**) The clustering separates the adductor muscle (green) from the venom gland (purple) based on the molecular pattern. (**C**) The inner gland structure is subdivided into a posterior (yellow), central (blue), and peripheral (red) sub-region (**SI Table S4**). (**A**) Spatial intensity map for a multivariate PCA within the full venom gland system. Highest variance (76%, PC1-3) is achieved for the first three principal components (PCs). (**D, E**) Principal component analysis (PCA) of respective tissue spots for the complete venom gland system and the various sub-regions (**Figure S4-5**).

Peak picking was applied to filter for relevant statistical signatures from the analyzed venom gland tissue sample, and resulted in 329 aligned peak features (*m/z* value range: 600–3200). An overlay of representative average spectra for respective muscle or venom gland regions highlighted highly associated peak features (**Figure S4**).

To allow a detailed correlation of mass spectral imaging data and morphological structures within the full venom gland system, we performed bisecting k-means clustering, an unsupervised multivariate segmentation analysis, to detect characteristic peak features clustered by correlation distance (**Figure 3**). The segmentation analysis revealed two distinct clusters of selected peak features and directly separated the closely associated AES muscle from the main elapid venom gland by specific molecular pattern (**Figure 3B**). Considering the specific nature and morphological structures of reptile muscles, which are complex organs made up of multinucleated long and cylindrical cells, called myofibrils, composed of typical myofilament proteins, like actin and myosin or other high-abundant muscle proteins, this clearly different segmentation is not unanticipated.^[37]^ Thus, we next performed a multivariate segmentation analysis solely on the venom gland and achieved a classification into three sub-regions along the venom gland based on the segmentation clusters (**Figure 3C**). Interestingly, unsupervised segmentation of spectral peak features showed a contrary segmentation to the aforementioned histological morphology (**Figure 3A**). While the histological morphology arranges the inner gland structure in anteroposterior regimes, peak features clearly show centrifugal segmentation (**Figure 3**). To confirm multivariate clustering of spatial peak features within the venom gland system, unsupervised principle component analysis (PCA), a dimensionality-reduction method, was applied to visualize the most discriminative principal components (PCs) for respective morphological regions (**Figure 3D-E** and **Figure S5**). While principal component 2 (PC-2, 25%) shows high discriminative power between the venom gland and the AES muscle due to specific peak features presumably associated to toxin peptides, PC-1 (40%) and PC-3 (11%) supported sub-areas within the venom gland beside the muscle region (**Figure S5**). Overall, peak features of muscle and gland tissue spots allow for a high variance (76%, PC1-3) in respective regions (**Figure 3D** and **Figure S5**). In addition, targeted PCA analysis (82%, PC1-3) specific to the venom gland supported a classification into three sub-regions along the venom gland (**Figure 3E** and **Figure S5**). Since the high variance can be associated to characteristic molecular signatures, in context to the morphological location, these findings demonstrate that both unsupervised and supervised multivariate methods support discriminative peptide toxin signatures within the venom gland system (**Figure 3** and **Figure S4-5**).

### Spatial venomics by MALDI-MSI

Despite the clustering of spectral and spatial peak features within the gland, we were particularly interested in identifying and spatially localizing various distinct toxin classes, named ‘spatial venomics’. To preserve spatial toxin resolution, tissue samples were fixed in formalin and embedded in paraffin in a biocompatible way to facilitate morphology preservation and MSI sample preparation. Alternative workflows include the application of other fixation solutions, i.e. ethanol-based fixative KinFix, as already describe for venom gland fixation.^[20,21]^ A clear advantage of ethanol-based fixation protocols is the possibility to measure intact toxins, which in turn is a disadvantage in terms of laborious downstream identification and inaccessibility of high molecular weight toxins. Further, immediate tissue fixation by shock freezing, which can be performed without sample embedding, presents an alternative option, though sample preparation and handling of frozen sections is difficult and leads to a decrease in peptide ion signals due to ionization suppression.

As a first step, we excluded all aligned peaks present in both, the muscle and venom gland region, to eliminate false-positives. The remaining 263 peaks (*m/z* value range: 600–3200) were aligned against our extensive peptide library thus permitting the identification of 175 peptide markers from 14 toxin families (**Table S5**). To cover various toxin classes within the entire venom gland system, we spatially localized them with at least two, and up to five, specific peptide markers. To verify the most discriminative toxin peptide marker, we performed statistical analysis by means of receiver operating characteristic (ROC) calculations for the different sub-regions, respectively. The area under the curve (AUC) can assume values between 0 and 1 and expresses the discrimination power of each toxin peptide (**Table S5**). The univariate procedure enabled identification of a large number of toxin families, such as 3FTx, 5NUC, CRISP, CVF, LAAO, NGF, PDE, PLA_2_, and svMP by a set of highly discriminative peptide masses (AUC >0.7 or <0.3), and thus enabled the various toxin classes to be spatially visualized within the different venom gland sub-regions (**Figure 4** and **Table S6**). In the central sub-region, which is oriented as a posteroanterior canal, the most discriminative peptide marker resulted from the toxin families PLA_2_, svMP, and CTx with high intensities in the anterior region (**Table S6**). The peripheral sub-region is partially associated with PDE toxins, although a spatial distribution map of specific PDE peptide markers shows a denser population in the posterior region of the venom gland. Interestingly, major toxin families are associated with high intensities in the posterior region of the venom gland, whereas none or low intensities of these toxin families were observed in the anterior parts of the venom gland (**Figure 4**). We detected a heterogeneous distribution of different toxin classes mainly along the posteroanterior canal with highly variable intensities within distinct regions. Some toxins were predominantly found in the posterior region (e.g., PDE and CRISP in **Figure 4**), others were found throughout the section but at higher levels in certain regions (e.g., PLA_2_, CTx, svMP in **Figure 4**), and some toxins exhibit a consistent distribution across the venom gland (e.g., 5NUC in **Figure 4**). The dominant abundance of 3FTx and the associated high number of peptide markers allowed us to spatially interrogate toxins at the level of various NTx sub-classes or CTx proteoforms (**Figure 5**). While the NTx toxin family shows a rather heterogeneous distribution, CTx can be uniformly detected throughout the venom gland via a combination of different specific peptide markers (**Figure 5A**). Visualization of localization at the proteoform level could only be accomplished by individual, specific peptide markers, which allowed accurate mapping and identified a heterogeneous distribution to distinct regions (**Figure 5B**). These analyses provide detailed insights into the fine distribution of different sub-classes of the 3FTx superfamily and show clearly delimited areas, e.g. sharp separation of lNTx and wNTx in the posterior region of the venom gland or the strictly heterogeneous distribution of different CTx proteoforms (**Figure 5B**). The identification and spatial localization of various sub-classes and proteoforms was achieved due to the very high abundance of this toxin family, though the combination of low abundance and reduced ionization potential preclude such a granular analysis for other toxin types.

**Figure 4.**
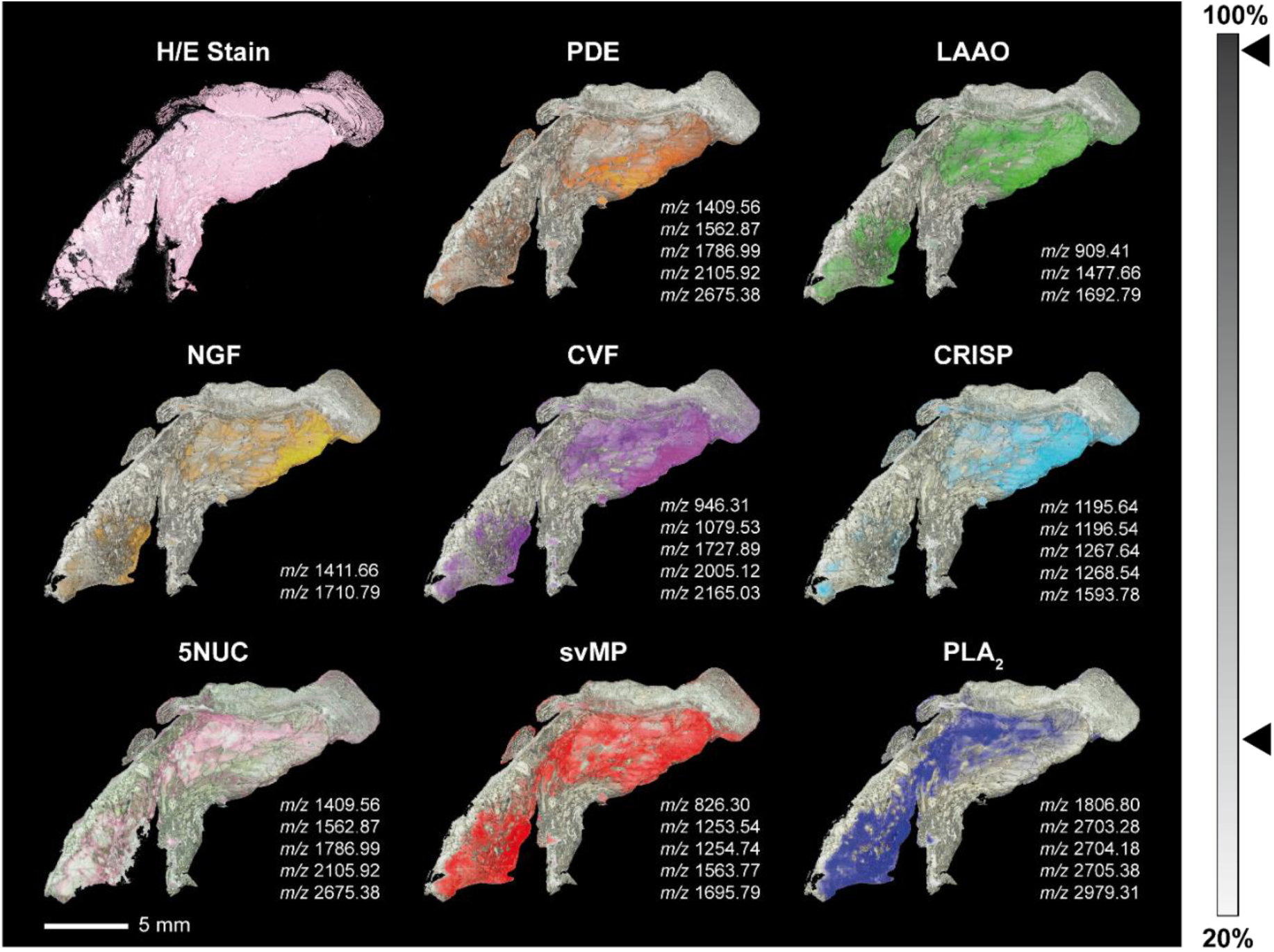
Spatial venom proteomics of midsagittal sections for the elapid venom system. Non-targeted MALDI imaging identifies different toxin families by heterogeneous peptide signatures within the venom gland system. The histological image was prepared after the MALDI imaging experiment (post-imaging) by H/E staining and is shown for tissue section orientation in MALDI-MSI analysis maps. The spatial segmentation of the venom gland system resulted in 263 extracted peak ions (±0.3 Da) from normalized spectra within an m/z 600-3200 mass range (**SI Table S3**). The spatial toxin distribution and relative intensities (grey scale bar) are shown for a set of characteristic MALDI m/z ion peaks for respective toxin families. The spatial visualization of different toxin families is based on the aforementioned color code (see **Figure 2**).

**Figure 5.**
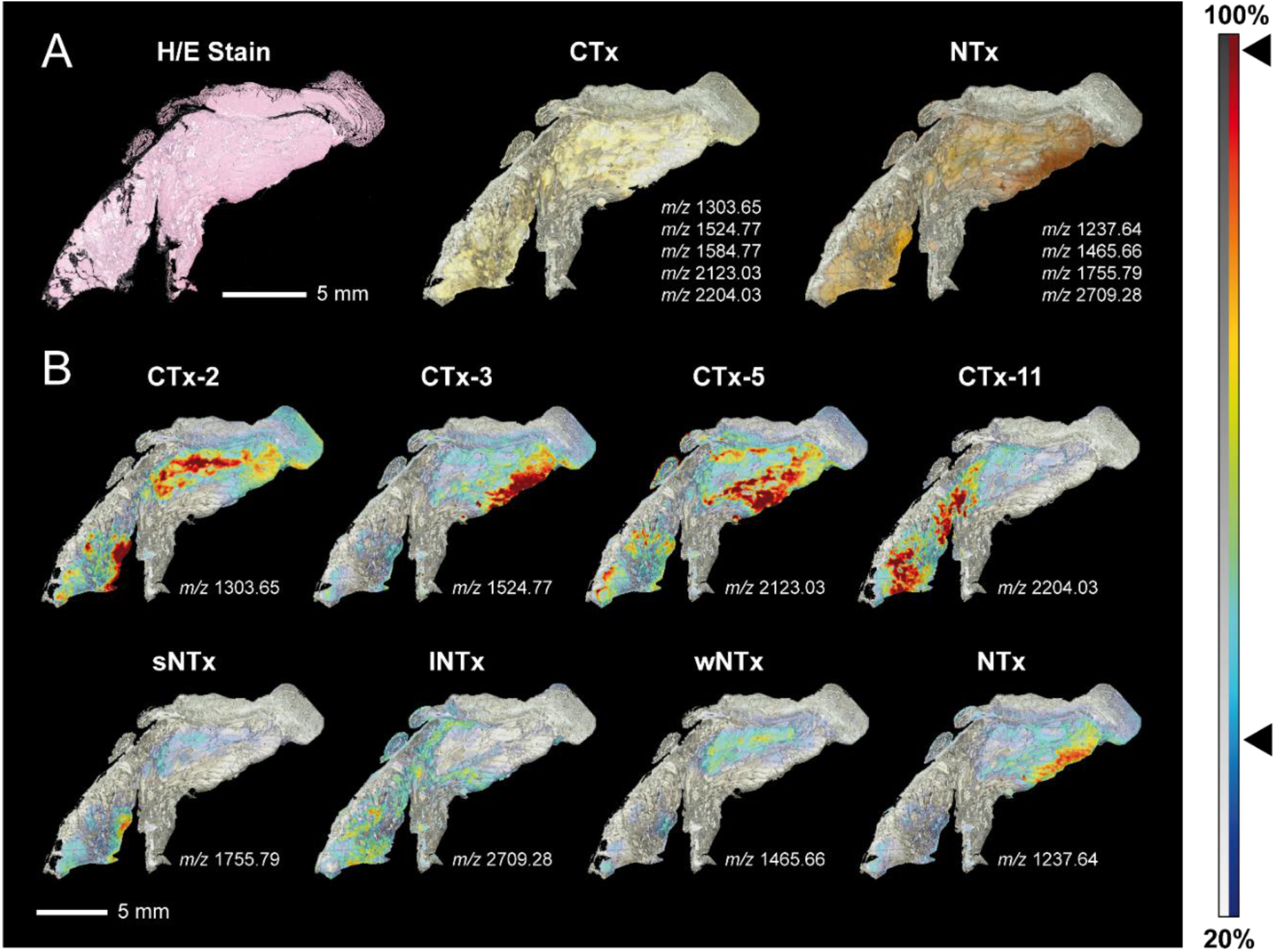
Spatial venom proteomics of 3FTx proteoforms and sub-groups. Non-targeted MALDI imaging allowed the identification of different neurotoxin subfamilies and cytotoxin proteoforms by heterogeneous peptide signatures within the venom gland system. The histological image was prepared after the MALDI imaging experiment (post-imaging) by H/E staining and is shown for tissue section orientation in MALDI-MSI spatial distribution maps. The spatial segmentation of the venom gland system resulted in 263 extracted peak ions (±0.3 Da) from normalized spectra within an m/z 600-3200 mass range (**SI Table S3**). The relative intensities (grey and color scale bar) are shown for characteristic MALDI m/z ion peaks for respective toxin families and proteoforms. (**A**) The spatial distribution and intensity for the neuro- (NTx) and cytotoxins (CTx) are shown for a set of characteristic MALDI m/z ion peaks. The spatial visualization of different toxin families is based on the aforementioned color code. (**B**) The heterogenous localization of various CTx proteoforms (upper line) and NTx subfamilies (bottom line) for specific peptide marker.

In summary, these results improve our knowledge of the basic biology of the snake venom gland as we demonstrate the identification and spatial localization of numerous toxin families in parallel. Our findings reveal that distinct toxin families, and functionally distinct sub-classes, exhibit spatial heterogeneity across the venom production system. To prevent proteolytic processes, spatial differentation is a simple and efficient tool. On the other hand, homogeneous distribution within the glandular tissue can help to increase antagonistic effects, such as inactivation of svMP by tripeptide inhibitors. We expect that these results will contribute to address the fundamental question as to how snake venom constituents are produced and stored within the venom glandular system.

## Conclusion

Herein MALDI-MSI was combined with a global venom analysis via proteotranscriptomics analyses to identify and spatially map various toxin families associated to the venom system of the Egyptian cobra (*Naja haje*). This particular approach allowed the identification of small and high-molecular toxin classes in near-cellular spatial resolution. Our findings highlight the molecular biology of centralized venom systems and the need for more holistic investigations of venom storage and delivery mechanisms. According to their morphological limitations and ecological context, the indirect venom modulation is of great importance to be considered in antivenom efficacy. Future studies on the multifunctionality of snake venoms can help to improve envenomation treatment. Thus, the simple venom-delivery system of an elapid with a complex venom arsenal might be a useful model system for gaining insight into the limitations placed on venom evolution by morphological constraints. Our results provide evidence for the venom optimization concept that toxins are non-uniformly abundant throughout simple venom systems of venomous animals, and that the link between ecology and toxin evolution may be more complex than previously assumed.

## Acknowledgements

In memory and thanks to Dr. Frank Mutschmann for the assistance of the gland removal. We would like to thank Marion Biering for the assistance of the venom gland preparation and Mrinalini for the assistance of the histological annotation. Furthermore, we thank Laura Ruysseveldt from the Herpetological Education & Research Project, Tierpark Berlin-Friedrichsfelde GmbH for providing the animal, and Klaus Rudloff for providing images of the living specimen. D.P. was supported by the German Research Foundation (DFG) through the CMFI Cluster of Excellence (EXC 2124).

