## Supplementary Information for "Spatial Venomics - Cobra Venom System Reveals Spatial Differentiation of Snake Toxins by Mass Spectrometry Imaging"

**Abstract:** Among venomous animals, toxic secretions have evolved as biochemical weapons associated with various highly specialized delivery systems on many occasions. Despite extensive research, there is still limited knowledge of the functional biology of most animal toxins, including their venom production and storage, as well as the morphological structures within sophisticated venom producing tissues that might underpin venom modulation. Here we report on the spatial exploration of a snake venom gland system by matrix-assisted laser desorption/ionization mass spectrometry imaging (MALDI-MSI), in combination with standard proteotranscriptomic approaches, to enable in situ toxin mapping in spatial intensity maps across a venom gland sourced from the Egyptian cobra (*Naja haje*). MALDI-MSI toxin visualization on the elapid venom gland reveals high spatial heterogeneity of different toxin classes at the proteoform level, which may be the result of physiological constraints on venom production and/or storage that reflects the potential for venom modulation under diverse stimuli.

**Keywords:** venom gland morphology • mass spectrometry imaging • spatial venomics • proteomics • venom heterogeneity

#### SUPPORTING INFORMATION — TABLE OF CONTENTS

**METHODS** 3

Specimen, venom collection and venom gland dissection 3

Computer tomography (CT) 3

Histology and Histochemistry 3

Venom gland transcriptome 3

Bottom-up venom proteomics 3

Top-down mass profiling 4

Mass spectrometry imaging (MSI) tissue preparation 4

MSI acquisition 4

MSI data analysis and visualization 5

MS data analysis 5

Data repository 5

**FIGURES** 5

Figure S1 6

Figure S2 6

Figure S3 7

Figure S4 7

Figure S5 8

**TABLES** 9

Table S1 9

Table S2 9

Table S3 9

Table S4 9

Table S5 9

Table S6 9

**REFERENCES** 10

#### SUPPORTING INFORMATION – METHODS

##### Specimen, venom collection and venom gland dissection

The venom was obtained from a single Morrocan male Egyptian *Naja haje* (*legionis*) individual, lyophilized and stored at +4 °C (**Fig. S1A**). The adult specimen of unknown age (>10 years old; snout to vent length of 164 cm, total length 194 cm) was kept at the Tierpark Berlin (Germany) since 2007 and euthanized in August 2017, due to tumorous occurrences. The healthy and intact venom glands were removed by veterinary supervision, snap frozen in liquid nitrogen, and stored at -80 °C until further processing (**SI Fig. S1B**).

##### Computer tomography (CT)

CT scans were performed with the 16 cm z-axis coverage and 0.5 mm resolution computer tomography system Aquilion ONE Genesis Edition (Canon Medical Systems, Zoetermeer, Netherlands) at the Leibniz-Institut für Zoo- und Wildtierforschung (IZW, Berlin, Germany). The postmortem *in situ* positioning on skeletal and soft tissue was documented using virtual coneXact double slice section technology with 640 slices every rotation (scan parameter: 120.0 kV, 200.0 mA, slice thickness 0.25 mm). Three-dimensional reconstructions were established by vitrea 2 software (Canon Vital Images, Veenendaal, Netherlands).

##### Histology and Histochemistry

Venom gland was fixed using 4% formalin in phosphate-buffered solution for 48 hours at 4 °C in the dark with a fixative volume five times of the tissue volume at least. The formalin-fixed tissue was placed in an embedding cassette and routinely processed by a fully-automated tissue processor Excelsior AS (Thermo Scientific, Bremen, Germany) by a solution series for stepwise dehydration and paraffin embedding. In short, seven runs (1 h; 30 °C) for dehydration were applied, starting with 50% ethanol (EtOH), followed by 70% EtOH, 80% EtOH and each two cycles with 96% EtOH and 99% EtOH. Then, three dehydration steps (0.5 h, 0.75 h, and 1 h; 30 °C) were performed with 100% xylene and treated with three cycles of paraffin wax (1 h, 2 h, 3 h; 62 °C) before the final embedding as paraffin blocks (Leica Biosystems, Nußloch, Germany). Venom gland sections were cut at 3 µm thickness using a microtome HM 340E (Thermo Scientific, Bremen, Germany) and directly placed on superfrost microscope glass slides (Thermo Scientific, Bremen, Germany). Subsequently, glass slides were incubated at least 3 h at 37 °C for adherence and dewaxed by a reversed solution series (10 min; RT), with the exception of two cycles 100% xylene and distilled water instead of 50% EtOH. Histological staining was performed with haematoxylin and eosin using standard protocols.[1]

##### Venom gland transcriptome

A separate specimen of *Naja haje*, from Uganda, was used for the construction of the venom gland transcriptome. The construction of the assembled transcriptome has previously been reported.[2,3] For annotation, assembled contigs were batch annotated with BLAST2GO Pro v3[4] using the BLASTx-fast algorithm with a significance threshold of 1e-5 against the NCBI non-redundant (NR) protein database. BLAST2GO annotations were inspected manually and preliminary annotation was performed by assigning contigs to toxins based on a combination of the BLAST2GO output and BLASTx searching sequences against the non-redundant protein database (nr). We built a species-specific assembled transcriptome database, extended by various *Naja* species[3] and the NCBI protein database (taxid: 8602).

##### Bottom-up venom proteomics

Proteomic analyses were guided by published protocols and are briefly summarized.[5–7] Briefly, crude venom was dissolved to a final concentration of 20 mg/mL (3% (v/v) acetonitrile and 1% (v/v) formic acid) and separated using a Supelco Discovery BIO wide Pore C18-3 column (4.6x150 mm, 3 µm particle size) coupled to an Agilent 1260 High-pressure Gradient System (Agilent, Waldbronn, Germany). The dried fractions from the aforementioned chromatographic separation were subsequently analyzed by precast SDS-PAGE (12% polyacrylamide with MES buffer, SurePAGE, GenScript, Piscataway Township, NJ, USA). Afterwards, the Coomassie-stained bands were excised and reduced with fresh dithiothreitol (100 mM DTT in 100 mM ammonium hydrogen carbonate, pH 8.3, for 30 min at 56°C) and alkylated with iodoacetamide (55 mM IAC in 100 mM ammonium hydrogen carbonate, pH 8.3, for 30 min at 25 °C in the dark). An in-gel trypsin (Thermo, Rockfeld, IL, USA) digestion was performed (6.7 ng/µL in 10 mM ammonium hydrogen carbonate with 10% (v/v) ACN, pH 8.3, for 18 h at 37°C, 0.27 µg/band). The peptides were extracted with 100 µL of aqueous 30% (v/v) ACN just as 5% (v/v) FA for 30min at 37°C. The supernatant was vacuum dried (ThermoSpeedVac, Bremen, Germany), redissolved in 30 µL of aqueous 3% (v/v) ACN with 1% (v/v) FA, and analyzed by LC−MS/MS analysis. The bottom-up analysis was performed with an Orbitrap XL mass spectrometer (Thermo, Bremen, Germany) via an Agilent 1260 HPLC system (Agilent Technologies, Waldbronn, Germany) using a reversed-phase Grace Vydac 218 MSC18 (2.1×150 mm, 5 µm particle size) column; with an injection volume of 20 µL and a flow rate of 0.3 mL/min. The chromatographic separation was performed with the following settings: after an isocratic equilibration (5% B) for 1 min, the peptides were eluted with a linear gradient of 5−40% B for 10 min and 40−99% B for 3 min, washed with 99%B for 3 min, and re-equilibrated in 5% B for 3 min. Further, complementary protein identification was performed with a specific peptide reference library recorded from the *N. haje* venom that was aligned to appropriate *m/z* values of the MSI measurements. Therefore, crude venom (50 µg) was digested in solution with or without the sample preparation kit (Biognosys, Schlieren, Switzerland) according to the manufacturer's protocol. The crude venom was dissolved in a dilution buffer (1 µg/µL), and a part mixed with 0.5% (v/v) reduction solution, and incubated for 30 min at 37 °C, 600 rpm. Afterwards, 16% (v/v) alkylation buffer was added and incubated at room temperature for 30 min in the dark. Next, sample was adjusted to a slightly basic pH (pH ⩾8), subsequently trypsin (2 µg) added to the solution, and incubated overnight at 37 °C, 600 rpm. In addition, we performed on-tissue trypsin digestion with paraffin removal, antigen retrieval, and tryptic digestion as described in detail for the MALDI-MSI tissue preparation below. Sample was acidified by adding 0.1% (v/v) TFA solution and centrifuged at 10,000 x g for 10 min. Afterwards, all samples underwent a C-18 ZipTip clean-up procedure (Millipore, Massachusetts, USA) and were submitted to nUPLC-MS/MS analysis using an analytical UPLC System (Thermo Dionex Ultimate 3000, Acclaim PepMap RSLC C18 column 75 µm x 15 cm; flow rate 400 nL/min, 70 min) coupled to an Impact II (QTOF-MS, Bruker Daltonik) in the data-dependent instant expertise mode.

**Top-down mass profiling**

Native and denaturing top-down proteomic experiments were performed as described previously and are briefly summarized.[6,7] In short, venom samples were dissolved in ultrapure water (10 mg/mL) and centrifuged at 12,000 x g for 5 min. Half of the dissolved sample (10 µL) was mixed with 10 µL of 0.5 M tris(2-carboxyethyl) phosphine (TCEP), and 30 µL of 0.1 M citrate buffer (pH 3) to reduce disulfide bonds. After 30 min incubation at 65 °C, samples were mixed with 50 µL of acetonitrile/formic acid/$H_2O$ (10:1:89, v/v/v), centrifuged at 12,000 x g for 5 min and 5 µL of supernatant of native and reduced sample injected for LC-MS/MS analyses. LC-MS/MS experiments of two technical replicates were carried out on a Vanquish ultra-high performance liquid chromatography (UHPLC) system coupled to a Q-Exactive hybrid quadrupole orbital ion trap (Thermo Fisher Scientific, Bremen, Germany). LC separation was performed on a Supelco Discovery Biowide C18 column (300 Å pore size, 2 x 150 mm column size, 3 µm particle size) using a flow rate of 0.3 mL/min.
Thermo data (.raw) were converted to a centroid mass spectrometry data format (.mzXML) using the MSconvert software of the ProteoWizard[8] package (http://proteowizard.sourceforge.net; version 3.0.10577): peak picking level 1+ and vendor prefer. The .mzXML data were deconvoluted to an .msalign file using TopFD (http://proteomics.informatics.iupui.edu/software/toppic/; version 1.3) with a maximum charge of 50, 100,000 Da maximum mass, MS1 S/N ratio of 3.0, MS2 S/N ratio of 1.0, *m/z* 3.0 precursor window and *m/z* 0.02 error. The final sequence annotation was performed by TopPIC[9] (http://proteomics.informatics.iupui.edu/software/toppic/; version 1.3) with the following settings: decoy database, 15 ppm mass error tolerance, e-value cutoff at 0.01 by E-value computation, 1.2 Da PrSM cluster error tolerance and a maximum of 2 mass shifts (±500 Da). The spectra were matched against an combined database of an in-house *Naja* transcriptome (532 entries), an *Naja* protein database (1834 entries, NCBI taxid 8638) and 116 cRAP protein sequences, manually validated and graphically visualized using the MS and MS/MS spectra by Qual Browser (Thermo Xcalibur 2.2 SP1.48).
The intact mass profile was inspected with the Qual Browser (Thermo Xcalibur 2.2 SP1.48) and Freestyle (Thermo Xcalibur 1.6.75.20). Deconvolution of isotopically resolved spectra was carried out by using the XTRACT algorithm of Thermo Xcalibur.

**Mass spectrometry imaging (MSI) tissue preparation**

Venom gland sections with 3 µm thickness were placed directly onto an indium-tin oxide (ITO) glass slide and heated at least 3 h at 37 °C for adherence. Afterwards, section slides were heated for 15 min at 80 °C and dewaxed by repetitive washes with two cycles xylene (100%, 5 min), one cycle isopropanol (100%, 5 min), decreasing concentration of ethanol (100%, 96%, 70%, 50%; each 5 min), and deionized water (5 sec) in order to improve sensitivity. Antigen retrieval (AR) was performed for 1 h at 60 °C in deionized water. On-tissue trypsin digestion and subsequent matrix overlay was performed using an automated spraying device (HTX TM-Sprayer, HTX Technologies LLC, Riemerling, Germany). The buffered trypsin solution (20 µg, 20 mM ammonium bicarbonate, 0.1% glycerol) was applied onto the section with a spray head temperature of 30 °C and a flow rate of 15 µL/min. After enzymatic incubation of tissue sections (2 h at 50 °C in a moiet chamber with a saturated potassium sulfate solution), matrix solution (7 g/L α-cyano-4-hydroxycinnamic acid in 50% acetonitrile and 1% trifluoroacetic acid) was overlaid using the aforementioned HTX Sprayer (75 °C, 120 µL/min).

**MSI acquisition**

MALDI-Imaging data acquisition was performed on a rapifleX MALDI Tissuetyper system (Bruker Daltonik GmbH, Bremen, Germany) operating in reflector mode, *m/z* 600–3200 detection range, 500 laser shots per spot, sampling rate of 1.25 GS/s, and raster width of 20 µm or 50 µm respectively. MSI was coordinated by FlexControl 3.0 and

FlexImaging 4.0 (Bruker Daltonik GmbH, Bremen, Germany) was used to establish the geometry and location of the section on the slide based upon the optical image and call upon FlexControl to acquire individual spectra, accumulating 200 shots per raster point. External calibration was performed using a peptide calibration standard (Bruker Daltonik GmbH, Bremen, Germany). After the MALDI-Imaging experiments, the matrix was removed with 70% ethanol and the tissue sections were stained with haematoxylin and eosin (HE) as histological overview staining.

**MSI data analysis and visualization**

Data analysis and visualization was performed using SCiLS Lab software (Version2015b, SCiLS GmbH, Bremen, Germany). MALDI-MSI raw data was imported into the SCiLS Lab software and converted to the SCiLS Lab file format and pre-processed by convolution baseline removal (width: 20) and total ion count (TIC) normalization. Segmentation pipelines as published previously were performed for regions of interest (ROIs) identification, specific tissue distinction, and peak finding. For external peptide library adjustment, *m/z* values from regions of interest (ROIs) were exported from SCiLS Lab SW as csv files.

**MS data analysis**

All raw LC-MS files were converted to open data format files (.mgf, .mzXML) via MSconvert GUI of the ProteoWizard[8] cross-platform package. The bottom-up analysis was performed with search engine PEAKS DB[10] (version 10.5, Bioinformatics Solutions Inc., Waterloo, Canada) using the following parameter settings: trypsin digestion, fixed modification: C (carbamidomethylation), variable modification: oxidation (M) and deamidation (N), precursor peptide tolerance: 10 ppm, peptide charge: 2+ to 8+, MS/MS tolerance: 0.8 Da, and maximum of one accepted miss cleavage. Identification of MALDI-IMS m/z values using the aforementioned peptide reference library requires the accordance of more than one peptide (mass differences ≤1.0 Da) to subsequently correctly assign the corresponding protein. Peptides with the lowest mass difference and highest -logP value compared to the LC-MS/MS reference list value were assumed to be a match in accordance with Cillero-Pastor et al. (2014) guidelines.[11]. The peptide assignment ('PAssT', https://github.com/benjaminhempel/PAssT) and multivariate analyses were performed with our in-house R scripts.

**Data repository**

The raw venom gland transcriptomic data has been deposited in the Sequence Read Archive (SRA) of NCBI (http://www.ncbi.nlm.nih.gov/sra) under accession SRR8206942, and the assembled contigs have been deposited in the Transcriptome Shotgun Assembly (TSA) database of NCBI (https://www.ncbi.nlm.nih.gov/genbank/tsa/) under accession GHRV00000000. All files are associated with the BioProject identifier PRJNA506018. Mass spectrometry proteomics data (.mgf, .raw, and .dat files) have been deposited with the ProteomeXchange Consortium[12] (http://proteomecentral.proteomexchange.org) via the MassIVE partner repository under project name "Spatial Venomics - Cobra Venom System Reveals Spatial Distinction of Snake Toxins by Mass Spectrometry Imaging" with the data set identifier PXD029870.

#### SUPPORTING INFORMATION – FIGURES

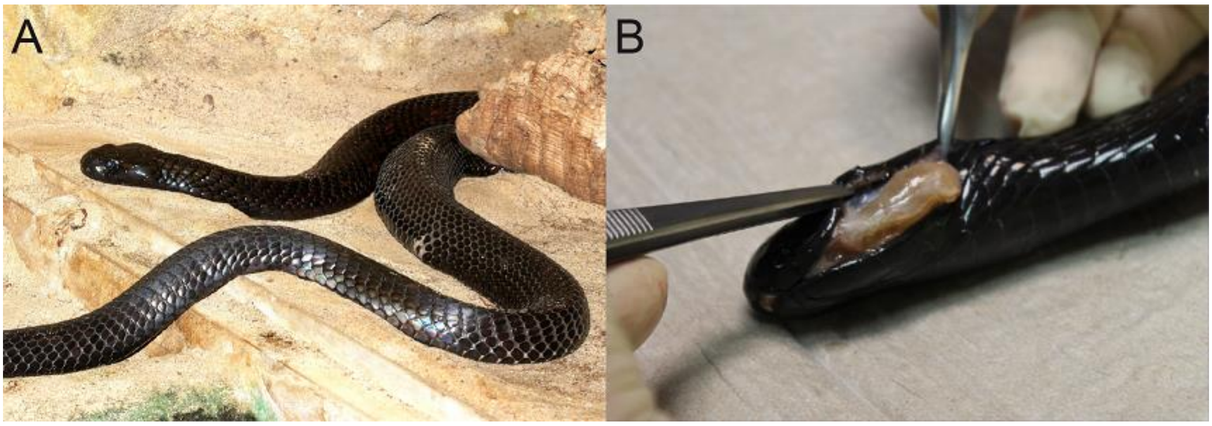


**Figure S1. Investigation of the Egyptian cobra (*Naja haje,* formerly *N. h. legionis*, locale: Morocco).** (**A**) Captive male specimen (>10 years old; snout to vent length of 164 cm, total length 194 cm) at the Tierpark Berlin (Germany, March 2017). The venom was obtained from a single Egyptian cobra, *Naja haje* (*legionis*), individual to perform proteomics analysis. (**B**) Intact venom glands were removed by veterinary supervision (August 2017). The elapid venom gland was fixed using 4% formalin in phosphate-buffered solution and subsequently paraffin-embedded.

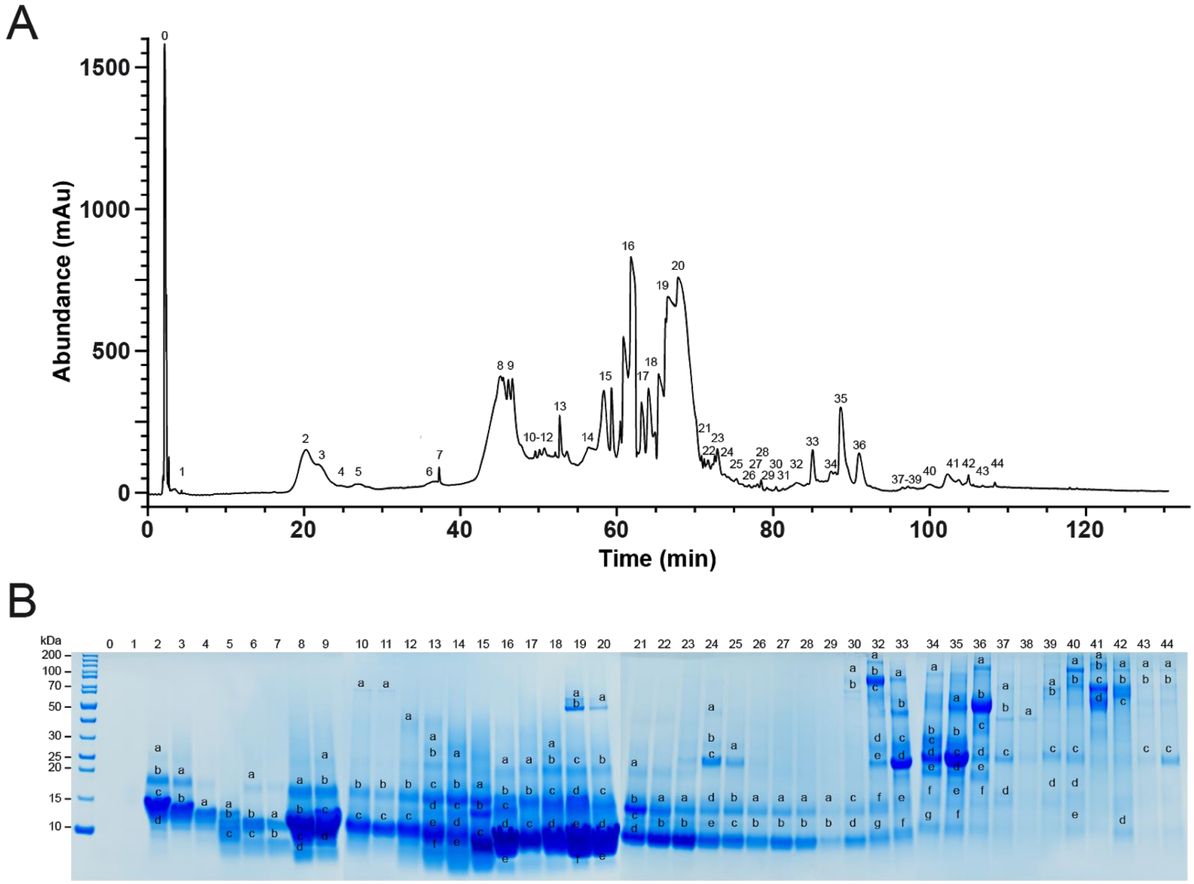


**Figure S2. Snake venomics workflow for bottom-up proteomics of *Naja haje* venom.** (**A**) RP-HPLC chromatogram performed by signal detection at UV absorbance λ = 214nm. Numbered fractions were subsequently submitted to 1D-SDS-PAGE separation. (**B**) The RP-HPLC fractions (indicated above the lane) of the elapid *N. haje* venom was analysed by SDS-PAGE under reducing conditions (Coomassie staining). Alphabetically marked bands per line were excised for subsequent tryptic in-gel digestion. A detailed nomenclature is shown in Table S2.

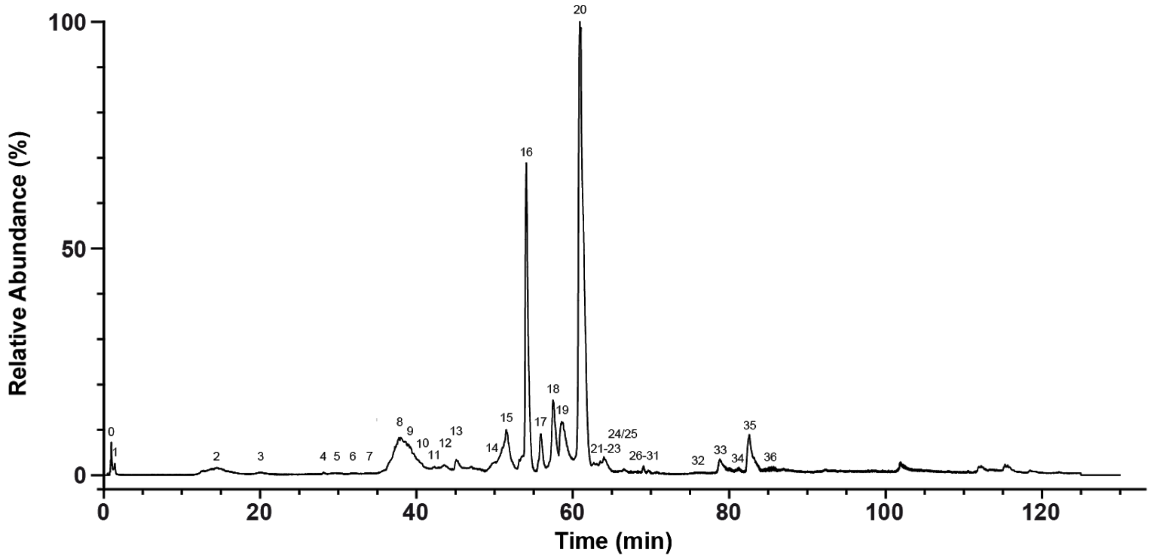


**Figure S3. Top-down mass profiling of *Naja haje* venom.** Total ion chromatogram (TIC) from native elapid *N. Haje* venom for top-down mass profiling. The total ion counts were measured by HPLC-ESI-MS and the relative abundance was set to 100% for the highest peak. Peak nomenclature based on HPLC peaks in Figure S2.

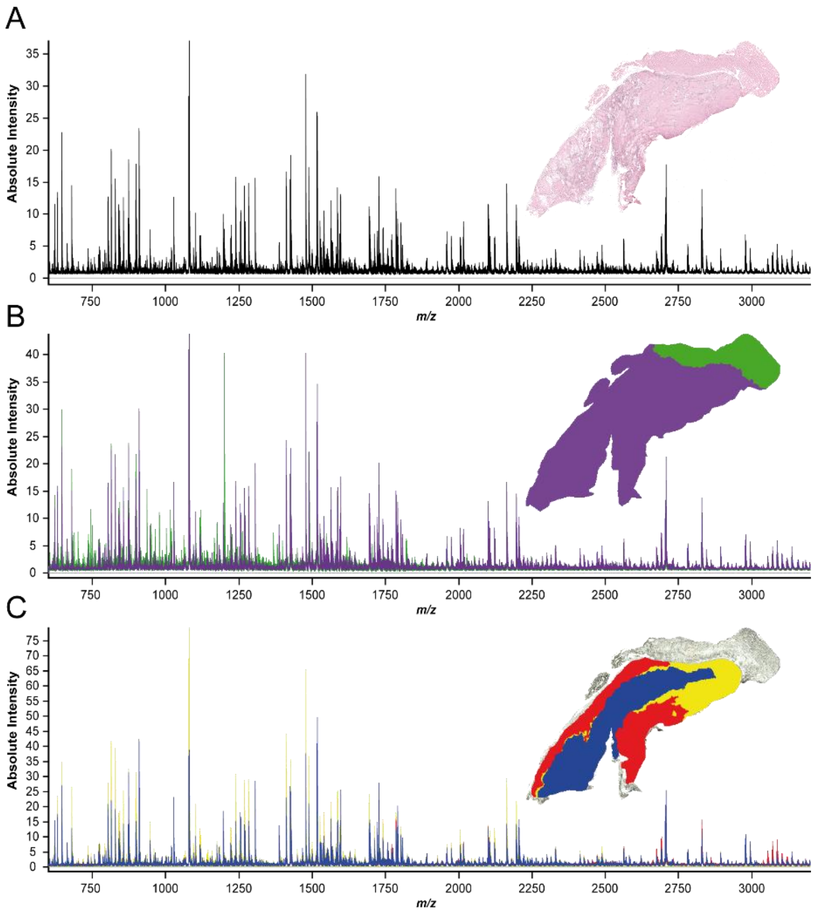


**Figure S4. Representative average spectra of venom system section.** Overview of average spectra for the full venom system (**A**) and overlay of average spectra for the venom gland vs. muscle (**B**) as well as all subregions within the venom gland (**C**).

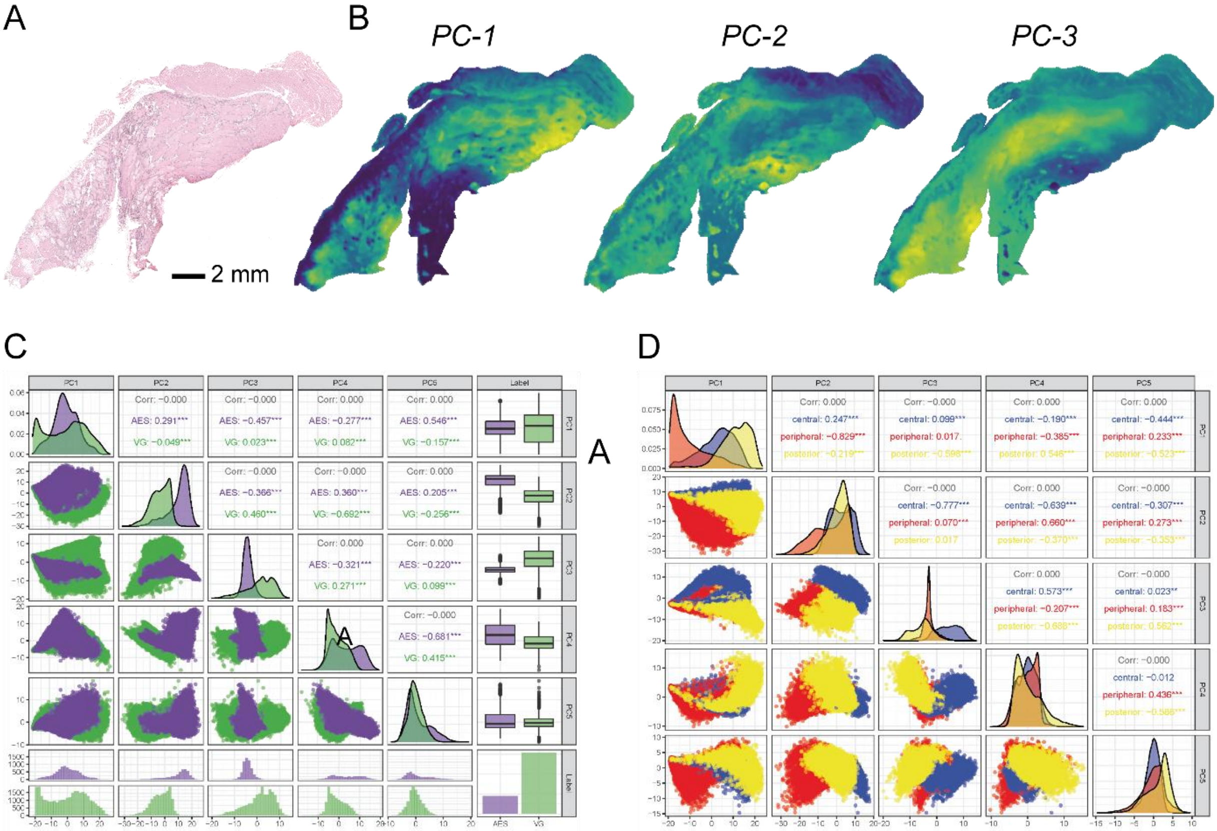


**Figure S5. Principal component analysis (PCA) of the venom gland system.** (**A**) The histochemical staining with hematoxylin and eosin (H&E) for tissue section orientation in segmentation maps of MALDI-MSI analysis. (**B**) Spatial intensity map for a multivariate PCA within the full venom gland system. Highest variance (76%, PC1-3) is achieved for the first three principal components (PCs). Variance of the peak features between the venom gland and muscle (AES) regions (**C**) as well as the different subregions (**D**).

### SUPPORTING INFORMATION – TABLES

**Table S1. The relative gene expression levels by venom gland transcriptomics from Egyptian cobra, *Naja haje*.** The construction of the assembled transcriptome has previously been reported.[2,3] List of full and partial length toxin transcripts with relative gene expression levels of the elapid toxin families are represented by a pie chart (**Figure 2A**). The corresponding bar chart reflects the relative gene expression levels of the secondary and minor categorized toxin families in the venom transcriptome (**Figure 2A**).

**Table S2. Shotgun proteomics analysis of the crude elapid venom.** Complementary protein identification was performed with a specific peptide reference library recorded by different shotgun LC-MS analysis workflows from crude *N. haje* venom. The crude venom was subjected to a trypsin digestion without (Method 1 &2, replicates) or with reduction and alkylation steps (Method 3), as well as an on-tissue digestion protocol (Method 4).

**Table S3. Alignment of venomic components by crude venom top-down mass profiling and bottom-up proteomics.** Fraction numbers are based on the RP-HPLC chromatogram (**Figure S2A**). Annotation was performed *de novo* and by peptide spectrum matching from in-gel digested protein bands (**Figure S2B**). Peak zero corresponds with injection peak Identification was carried out against a species-specific assembled transcriptome database, extended by various *Naja* species[3], the NCBI protein database (taxid: 8602) and a set of proteins found as common contaminants (cRAP). SDS-PAGE and top-down mass profile analysis provided the average molecular weight. Intact mass calculation was performed by isotopic resolved deconvolution with Thermo Xtract.

**Table S4. Relative venom proteome quantification.** The quantification of venom composition is based on the RP-HPLC peak integration ($UV_{214nm}$) in comparison to the total integral of all analyzed peaks. The relative venom protein abundance (proteome) of the major 3FTx toxin family and subgroups, secondary (>5%) as well as minor (>0.5%) toxin families, are represented by a pie chart (**Figure 2B**). The corresponding bar chart reflects the relative protein abundance of the secondary and minor categorized toxin families in the venom proteome (**Figure 2B**).

**Table S5. Peptide mass assignment of MSI data.** Identification of MALDI-MSI m/z values was performed with our aforementioned peptide reference library (**Table S2**). Peptides with the lowest mass difference and highest -logP value compared to the LC-MS/MS reference list value were assumed to be a match (<1.0 Da) in accordance with Cillero-Pastor et al. (2014) guidelines.[11].The peptide assignment tool and multivariate analyses were performed with our in-house R script.

**Table S6. Spatial distribution of toxin families within the venom gland system**. Strong (▲), moderate (●) and weak (Δ) abundances are indicated for different regions. Within the rostral venom gland (RVG), caudal venom gland (CVG), dorsal cap (DC) and ventral base (VB) and observed anterior (ant.), inferior (inf.), medialis (med.) and posterior (post.) subregions.

### SUPPORTING INFORMATION – REFERENCES
